# *singleCellHaystack*: A clustering-independent method for finding differentially expressed genes in single-cell transcriptome data

**DOI:** 10.1101/557967

**Authors:** Alexis Vandenbon, Diego Diez

## Abstract

**Summary:** A common analysis of single-cell sequencing data includes dimensionality reduction using t-SNE or UMAP, clustering of cells, and identifying differentially expressed genes. How cell clusters are defined has important consequences in the interpretation of results and downstream analyses, but is often not straightforward. To address this difficulty, we present a new approach called *singleCellHaystack* that enables the identification of differentially expressed genes (DEGs) without relying on explicit clustering of cells. Our method uses Kullback-Leibler Divergence to find genes that are expressed in subsets of cells that are non-randomly positioned in a multi-dimensional space. We illustrate the usage of *singleCellHaystack* through applications on several single-cell datasets, demonstrate that it enables the identification of markers important for cell subset separation in an unbiased way, and compare its results with those of a traditional, clustering-based DEG prediction method.

**Availability and implementation:** *singleCellHaystack* is implemented as an R package and is available from https://github.com/alexisvdb/singleCellHaystack

## 1 Introduction

Recent advances in single-cell technologies enable us to assess the state of cells by measuring different modalities like RNA and protein expression with single cell resolution (1–5). Since the appearance of the first single-cell technologies, hundreds of bioinformatics tools have been developed to process, analyze and interpret the results from single cell omics data (6), such as Monocle2 and Seurat (7, 8).

A standard protocol for analyzing single-cell data includes dimensionality reduction methods, such as PCA, t-distributed stochastic neighbor embedding (t-SNE) and Uniform Manifold Approximation and Projection (UMAP) to represent the data in fewer (typically 2) dimensions (9, 10). Finally, cells are clustered, and differentially expressed genes between the different clusters are identified. This approach for finding differentially expressed genes by comparing between clusters is widely used in existing methods (8, 11, 12), and enables finding cluster-specific marker genes that facilitate labeling different cell populations. However, recent comparisons found that DEG prediction approaches for bulk RNA-seq do not generally perform worse than methods designed specifically for single-cell RNA-seq (scRNA-seq), and that agreement between existing methods is low (13, 14). Defining more flexible statistical frameworks for predicting complex patterns of differential expression is one of the grand challenges in single cell data analysis (15).

One problem with clustering-based approaches for DEG prediction is that the definition of cell clusters is often not straightforward. The number of biologically relevant clusters naturally occurring in a dataset is often not obvious. The high-dimensionality of the data makes it hard to evaluate if the number of clusters and their borders make sense or if they are arbitrary. Furthermore, some cell sub-populations may not be clustered independently and their defining signature may end up being obscured within a larger cluster. This can be critically important for low abundance populations in experiments using unsorted cells from tissue, where only a few representative cells may be present. Thus, the clustering of cells has important consequences in the interpretation of results and downstream analyses.

To address this problem we present *singleCellHaystack*, a methodology that uses Kullback-Leibler Divergence (*D*_*KL*_; also called relative entropy) to find genes that are expressed in subsets of cells that are non-randomly positioned in a multi-dimensional (≥2D) space (16). In our approach, the distribution of cells expressing or not expressing each gene is compared to a reference distribution of all cells in the input space. From this, the *D*_*KL*_ of each gene is calculated, and compared with randomized data to evaluate its significance. Thus, *singleCellHaystack* does not rely on clustering of cells, and can identify differentially expressed genes in an unbiased way.

An R package for running *singleCellHaystack* analysis and additional functions for visualization and clustering of genes is available at https://github.com/alexisvdb/singleCellHaystack.

## 2 Materials and methods

Our approach contains two main functions: `haystack_2D` and `haystack_highD`, for 2D and multi-dimensional (≥2D) input spaces, respectively. The concept of both functions is the same, and is briefly explained below. More details are given in Supplementary Material. We refer to Supplementary Fig. S1 for an overview of the workflow.

### 2.1 *singleCellHaystack* methodology

*singleCellHaystack* uses *D*_*KL*_ to estimate the difference between a reference distribution of all cells in a multi-dimensional space (distribution *Q*) and the distributions of the cells in which a gene *G* was detected (distribution *P*(*G* = *T*)) and not detected (distribution *P*(*G* = *F*)).

For 2D spaces (such as typical t-SNE or UMAP plots), `haystack_2D` divides the 2D space into a grid along both axes. For multi-dimensional spaces (such as the first several principal components), `haystack_highD` defines a set of grid points covering the subspace in which the cells are located (see Supplementary Material).

Next, a Gaussian kernel is used to estimate the density of cells at each grid point. Summing the contributions of all cells gives us *Q*; the subset of cells in which *G* is detected gives us *P*(*G* = *T*); and the subset of cells in which *G* was not detected gives us *P*(*G* = *F*). Each distribution is normalized to sum to 1.

The divergence of gene *G*, *D*_*KL*_(*G*), is calculated as follows:

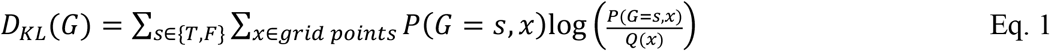

where *P*(*G* = *s*, *x*) and *Q*(*x*) are the values of *P*(*G* = *s*) and *Q* at grid point *x*, respectively.

Finally, the significance of *D*_*KL*_(*G*) is evaluated using randomizations, in which the expression levels of *G* are randomly shuffled over all cells. The mean and standard deviation of *D*_*KL*_(*G*) in randomized datasets follow a clear pattern in function of the number of cells in which a gene was detected (see Supplementary Fig. S2 for examples), which is modeled using B-splines (17). P-values are calculated by comparing the observed *D*_*KL*_(*G*) to the predicted mean and standard deviation (log values).

### 2.2 *singleCellHaystack* advanced options

The distribution *Q* and the randomizations described above ignore the fact that some cells have more detected genes than others. *singleCellHaystack* can be run in an advanced mode, in which both the calculation of *Q* and the randomizations are done by weighting cells by their number of detected genes (see Supplementary Material for more details).

In addition, *singleCellHaystack* includes functions for visualization and clustering gene expression patterns in the multi-dimensional space.

### 2.3 scRNA-seq datasets and processing

We downloaded processed data (read counts or unique molecular identifiers) of the Tabula Muris project (FACS-sorted cells: 20 sets; Microfluidic droplets: 28 sets), the Mouse Cell Atlas (Microwell-seq: 87 sets) and a dataset of several hematopoietic progenitor cell types (5, 18, 19). For each dataset, cells and genes were filtered, the 1,000 most variable genes were selected, and dimensionality reduced using PCA. Subsequently, the first 50 Principal Components (PCs) were used as input for t-SNE and UMAP analysis, following the recommendations by Kobak and Berens (20). Finally, *singleCellHaystack* was run on both 2D (2D t-SNE and UMAP coordinates) and multi-dimensional (5, 10, 15, 25, or 50 PCs) representations of each dataset to find genes with biased expression patterns.

### 2.4 Known cell type marker genes

We downloaded marker gene data from the CellMarker database (21). A total of 7,852 unique mouse gene symbols are included as marker genes, which we split into 630 “high-confidence” markers (reported in ≥5 publications), and 7,222 “low-confidence” markers (reported in 1 to 4 publications). Other genes we regarded as “non-marker” genes.

### 2.5 Generating a simulated dataset

We generated an artificial single-cell dataset using Splatter (22). This artificial data contained 10,000 genes in 2,000 cells divided into 5 “cell types” (using the method=“groups” setting in Splatter). Otherwise default parameters were used. After filtering out genes detected in less than 50, or more than 1,950 cells, 7,585 genes remained. Read counts were processed to counts per million in each cell, and the PCA was performed on the log-transformed data. The first 5 PCs were used as input for `haystack_highD`.

The output of Splatter contains differential expression factors (“DEFac[Group]”), showing whether a gene has differential expression (factor different from 1) or not (factor = 1) in each group. We defined a “differential expression score” for each gene as the sum of the log_2_-transformed factors (Supplementary Fig. S3). We regarded the 1,857 genes with scores > 0.3 as DEGs. A precision-recall curve was made using the ROCR package (version 1.0-7) in R (23).

To the Splatter dataset we manually added a gene with a biased expression pattern that is independent of the 5 cell types. This gene had no expression in any cells, except in the 200 cells that are closest to cell No. 300 in the space defined by the 1^st^ and 2^nd^ PC (Supplementary Fig. S4).

### 2.6 Predicting DEGs using Seurat’s *FindAllMarkers* function

To compare runtimes and results of *singleCellHaystack* with those of Seurat (version 3.1.0), we used the same 50 PCs of each dataset to define clusters of cells in the data using default options of the functions *FindNeighbors* and *FindClusters* (using the default Louvain algorithm). Next, DEGs were predicted between the resulting clusters of cells using the *FindAllMarkers* function. We used default options, except for options “only.pos = FALSE” and “return.thresh = 1” in order to obtain results that were comparable to those of *singleCellHaystack*. The default test used in *FindAllMarkers* is the Wilcoxon Rank Sum test. Runtimes of the *FinallMarkers* function were compared to those of our method.

The above workflow results in a several p-values for every gene (1 p-value for the comparison of every cluster with all other clusters). For every gene, we retained the minimum p-value, and compared those to the p-values returned by *singleCellHaystack*.

## 3 Results and Discussion

### 3.1 Application on simulated data

To illustrate the validity of our approach, we applied *singleCellHaystack* on a simulated dataset containing 2,000 cells in 5 “groups” (i.e. simulated cell types; Fig. 1A). The inputs to our method were the coordinates of the 2,000 cells in the first five PCs, and the detection levels of 7,585 genes in these cells. *singleCellHaystack* successfully predicted DEGs (Fig. 1B-E), even though it works independently of cell clustering, and we did not provide it with the cell group data. The five top biased genes predicted by our method had clear non-random expression patterns (Fig. 1B). To get a more global view on the performance of *singleCellHaystack* we calculated a “differential expression score” for every gene based on the Splatter simulation data (see Materials and methods), reflecting the gene’s strength of differential expression. We confirmed that *singleCellHaystack* gave high scores to genes with strong biases in expression between the five cell groups (Fig. 1C), and that it could predict DEGs with high precision (Fig. 1D).

**Figure 1:**
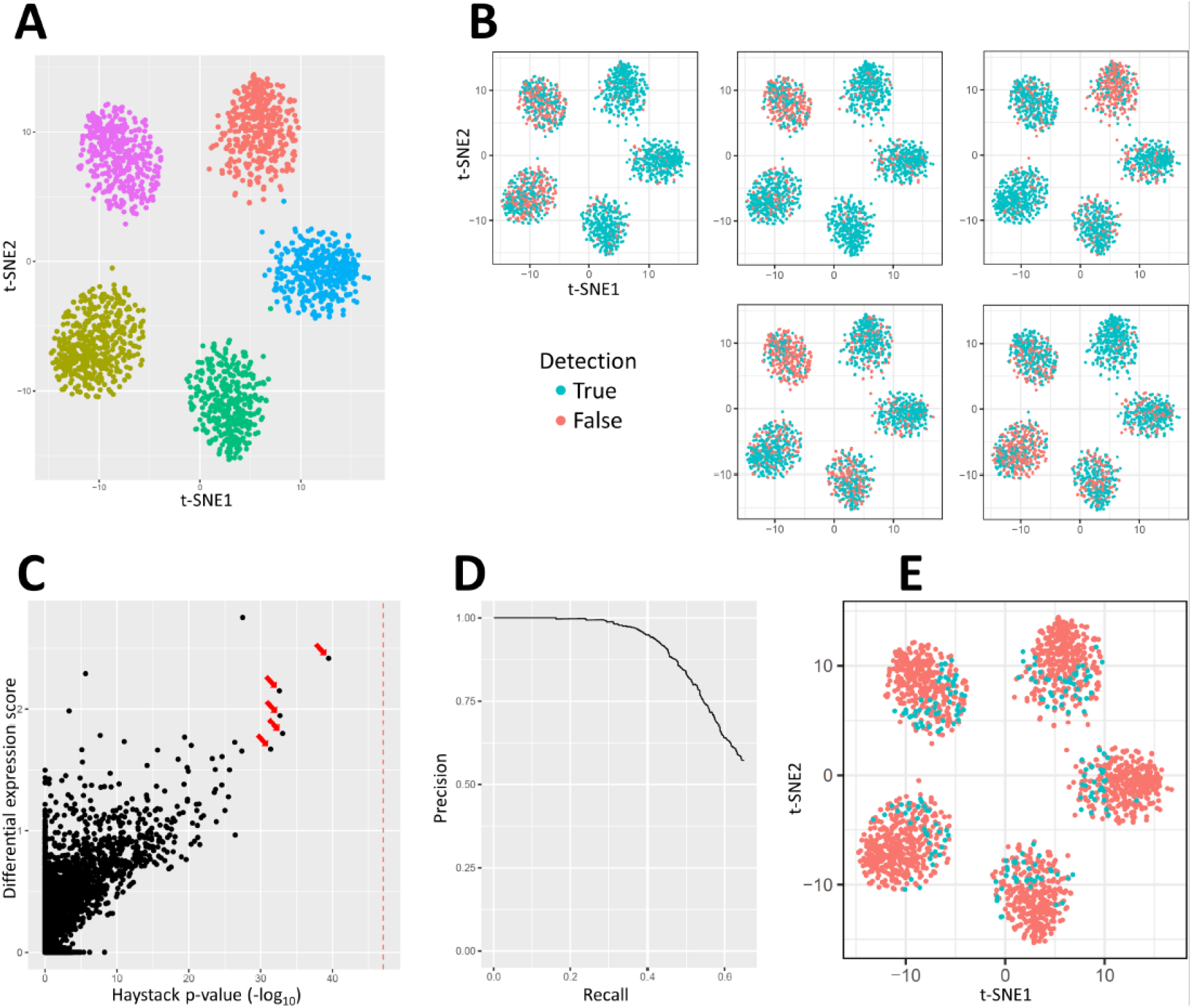
Application of *singleCellHaystack* on an artificial dataset. **(A)** t-SNE plot of the artificial dataset. Colors indicate cell groups **(B)** t-SNE plots for the five top-scoring genes predicted by *singleCellHaystack*. **(C)** Scatterplot of p-values of our approach (X-axis) and differential expression scores (Y-axis) of all genes. The five top-scoring genes of **(B)** are indicated by arrows. The dotted line represents the p-value of the manually added gene shown in **(E)**. **(D)** Precision-recall curve of our method on this artificial dataset. Positives were defined as genes with differential expression score > 0.3. **(E)** t-SNE plot of a manually constructed gene with no differential expression between cell groups, but with a strongly biased expression pattern in the first and second PC.

It should be noted that differential expression in the Splatter simulation data is completely group-based (i.e. there are no DE patterns that go beyond the cell grouping). To further illustrate that *singleCellHaystack* does not rely on grouping or clustering of cells, we manually added a gene with highly biased expression in the plane defined by the first and second PC, independent of the cell clusters (see Materials and methods, Fig. 1E and Supplementary Fig. S4). This highly biased DEG was correctly detected by *singleCellHaystack*, with a p-value of 8.7e-48 (red dotted line in Fig. 1C). This gene is not detected by methods relying on comparison between clusters of cells.

These results show that *singleCellHaystack* can be used to detect DEGs in real single-cell datasets, where cell groupings are often not obvious, without the need to group cells into arbitrary clusters.

### 3.2 Application on real single-cell datasets

We applied *singleCellHaystack* on 136 real scRNA-seq datasets of varying sizes (149 to 19,693 cells). Median runtimes of `haystack_highD` with 50 PC inputs were and 102 and 115 seconds using the simple and advanced mode, respectively. Runtimes followed an approximately linear function of the number of cells in each dataset (Supplementary Fig. S5A-B). Runtimes for 5, 10, 15, or 25 PC input were similar (not shown). Median runtimes for `haystack_2D` on 2D t-SNE coordinates were 75 and 84 seconds using the simple and advanced mode, respectively (Supplementary Fig. S5C-D).

In all datasets, large numbers of genes were found to have significantly biased distributions in the input spaces. This observation in itself is not surprising, since samples typically include a variety of different cell types, often forming loose clusters in the PC space. Rather than interpreting *singleCellHaystack* p-values in the conventional definition, the ranking of genes is more relevant.

As an illustration of the usage of *singleCellHaystack*, we here present 3 example results of datasets based on different sequencing technologies. In all three cases, the coordinates of cells in the first 50 PCs was used as input, along with the detection levels of all genes in all cells.

Figure 2 summarizes the result of the Tabula Muris marrow tissue dataset (FACS-sorted data). 5,250 cells and 13,756 genes were used as input, and the *singleCellHaystack* run took 225s in the default mode. The t-SNE plot shows a typical mixture of clearly separated as well as loosely connected groups of cells, with considerable variety in the number of genes detected (Fig. 2A). The gene with the most significantly biased expression was *Stmn1*, which is detected only in a subsets of cells (Fig. 2B). To illustrate the variety in expression patterns, we grouped biased genes into 5 clusters based on hierarchical clustering of their expression in the 50 PC input space (Supplementary Fig. S6). Fig. 2C-F show the most significantly biased genes of the other 4 groups.

**Figure 2:**
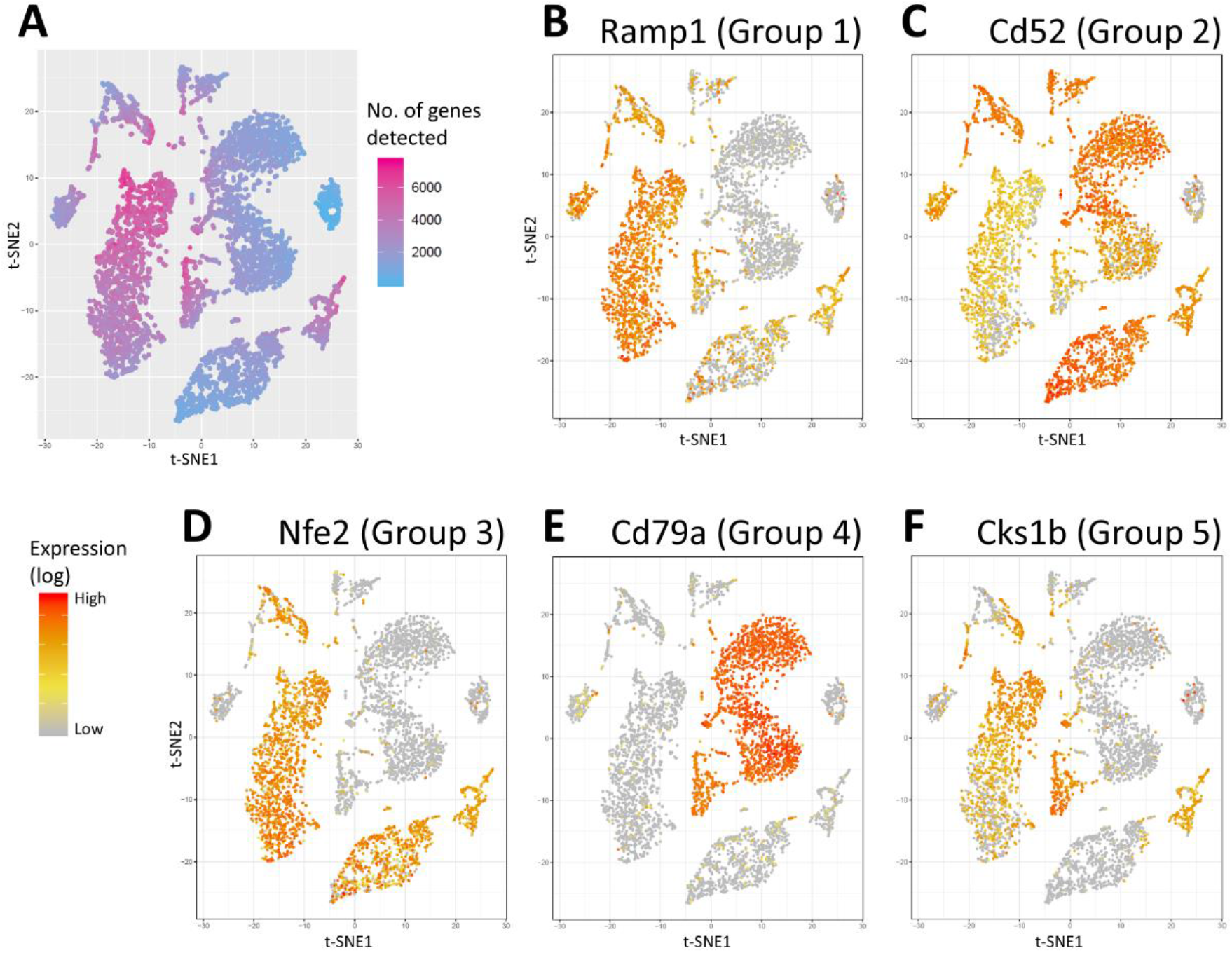
Application of *singleCellHaystack* on marrow tissue dataset. **(A)** t-SNE plot of the 5,250 cells. The color scale shows the number of genes detected in each cell. **(B-F)** Expression patterns of five highly biased genes, representative of the five groups in which the genes were clustered.

Results for two other example datasets are shown in Supplementary Fig. S7 and S8.

### 3.3 Known cell type marker genes often have biased expression patterns

Focusing on results of `haystack_highD` applied on the first 50 PCs of each dataset, we investigated whether genes with strongly biased expression are often known marker genes. For each dataset, we ranked genes by their p-value, and counted how often the genes at each rank were high-confidence markers, low-confidence markers, or non-marker genes (see Materials and methods). High-confidence marker genes (such as *Cd45* and *Kit*) were strongly enriched among top biased genes: although they comprise only 2.2% of all genes, on average 32.4% of the top 50 ranked genes were high-confidence cell type markers (Fig. 3).

**Figure 3:**
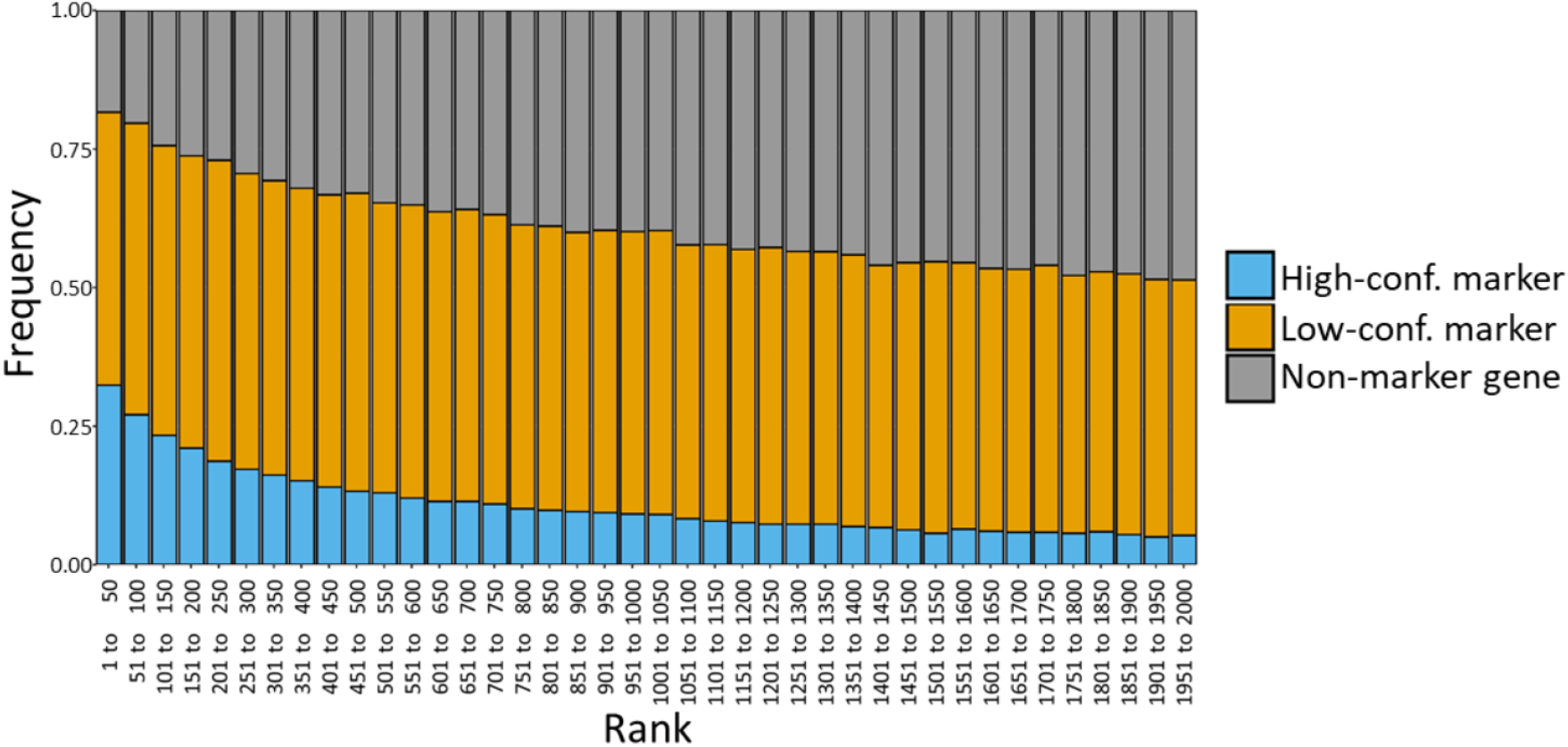
Frequencies of markers genes among genes with biased expression patterns. The frequencies of high-confidence (blue), low-confidence (orange), and non-marker genes (grey) among genes with biased expression in all datasets. The X-axis shows ranks in bins of 50 (ranks 1 to 50, ranks 51 to 100, etc).

### 3.4 The advanced mode takes into account general gene detection levels

In scRNA-seq data, there can be considerable variation in the number of detected genes in each cell. In some datasets this results in clusters of cells with higher or lower general detection levels. The “advanced” mode of *singleCellHaystack* can be used to find genes that have biased expression patterns that are contrary to the general pattern of detected genes. Figure 4 shows three example results, comparing the “advanced” mode to the “default” mode. The top biased gene in the “advanced” mode is often expressed in cells that have in general fewer detected genes.

**Figure 4:**
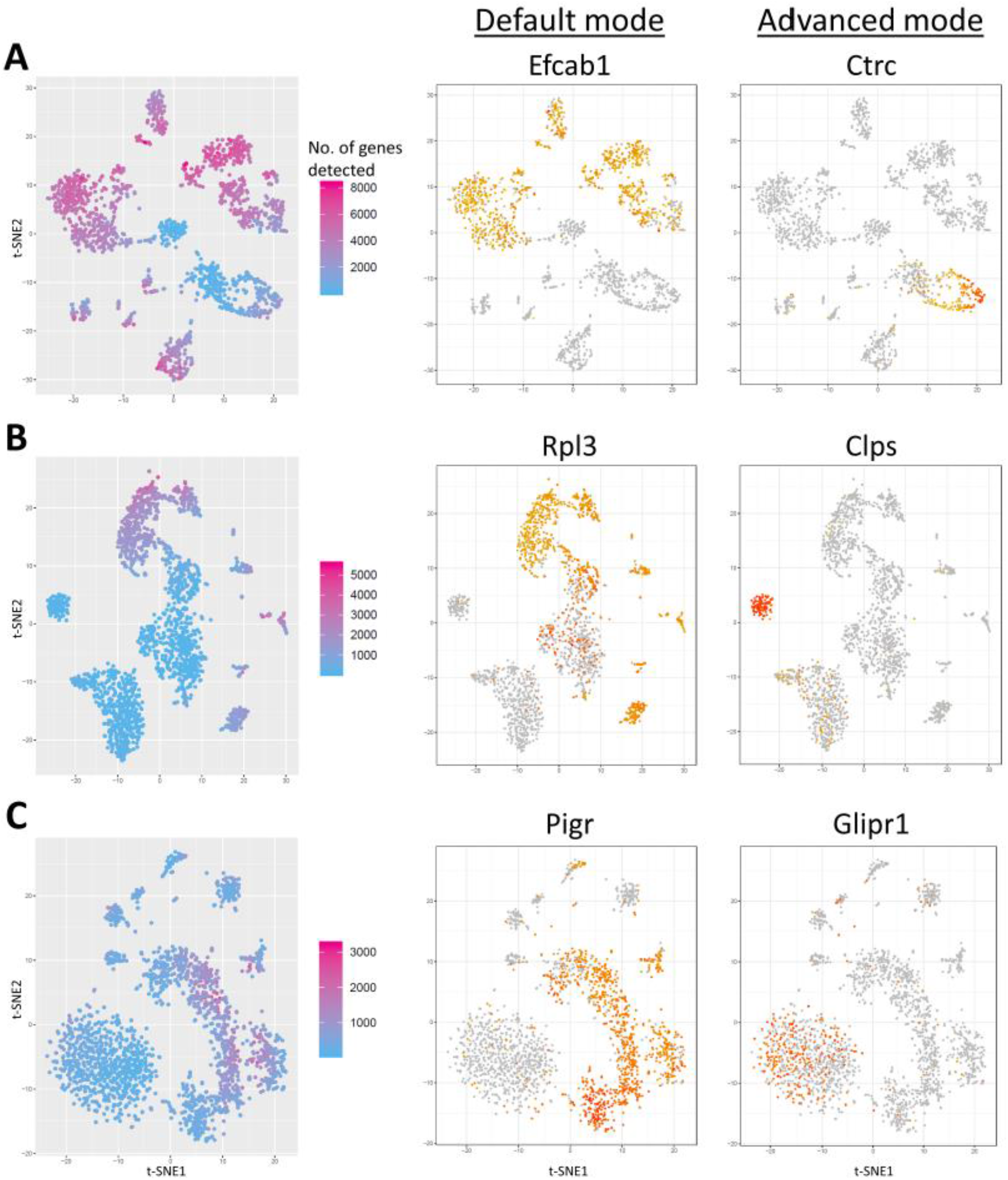
Example results of “default” versus “advanced” mode of *singleCellHaystack*. A t-SNE plot (left), the most strongly biased gene in the default mode (center) and advanced mode (right) are shown for **(A)** the Tabula Muris pancreas (FACS-sorted data) **(B)** the Tabula Muris lung (P8 12; Microfluidic droplet) and **(C)** the Mouse Cell Atlas small intestine 2 dataset.

### 3.5 Consistency of results

The definition of the grid points in `haystack_highD` is not deterministic (see Supplementary Material). As a result, grid points differ between each run. To evaluate how much this impacts results, we ran `haystack_highD` ten times on each dataset using the first 5 PCs as input. For each dataset, we calculated the mean rank of each gene in the results and compared the ranking in individual runs to the mean ranking. In general, there was a high consistency in the ranking of genes, suggesting that the differences in grid points have only a limited impact on results (Supplementary Fig. S9A).

In addition, we observed that the results of applications on 5 PCs were in general consistent with applications of `haystack_2D` on t-SNE and UMAP coordinates (both based on 50 PCs, Supplementary Fig. S9B-C), confirming that part of the information contained in the first 50 PCs is indeed captured in the t-SNE and UMAP coordinates. In contrast, there was somewhat larger discrepancy with results using only the 2 first PC coordinates as input (Supplementary Fig. S9D). Discrepancies increased as the difference in input dimensionality increases (10 PCs < 15 PCs < 25 PCs < 50 PCs; Supplementary Fig. S9E-H).

### 3.6 Comparison with other methods

Several computational methods for predicting DEGs exist (for example: 8, 11, 12), all relying on a comparison between predefined groups of cells. When researchers have a good prior knowledge of the cell types that are present in their sample, they can use the expression of marker genes (RNA or protein level) to define groups of cells, and predict DEGs by comparing between groups.

More typically, the subpopulations of cells in the data are not well known, and exploratory analysis is needed. In this case, a typical workflow starts with the definition of groups of cells, using for example clustering by the Louvain or Leiden algorithms (24, 25). In a typical dataset of thousands or ten thousands of cells, this results in between 5 to >20 clusters of cells. Subsequently, DEGs are usually predicted by comparing each individual cluster to all other clusters together, thus biasing DEGs towards genes that are expression exclusively in one or a few clusters. Naturally, a comprehensive prediction of DEGs between combinations of clusters is not practical or even feasible (ex: there are 190 pairs and 1,140 triplets of clusters in 20 clusters). On top of that, there is an inherent difficulty in deciding a suitable number of clusters in data that is high-dimensional and therefore hard to visualize. Finally, consistency between DEG prediction methods has been reported to be low (14). In contrast, *singleCellHaystack* works independently of any grouping or clustering of cells.

Here, we present a comparison between our method and the default test used in Seurat’s *FindAllMarkers* function (Wilcoxon Rank Sum test), one of the most widely used approaches. The median runtime of the Wilcoxon Rank Sum test on our 136 datasets was 452 seconds, which is about 4 times slower than our method (paired t-test p-value 8.9e-10). *singleCellHaystack* runtimes were shorter in 125 out of the 136 datasets. The *FindAllMarkers* runtimes did not depend so much on the number of cells in dataset (Fig. S5E), but rather on the number of predicted clusters of cells (Fig. S5F).

As a representative case, we show the comparison between *singleCellHaystack* and *FindAllMarkers* on the Tabula Muris marrow dataset (Fig. 5). In general, the agreement between both methods was low (Fig. 5 top-left, Fig. S10). The top 100 high-scoring genes of both approaches have only 3 genes in common (Fig. S10). To gain understanding into the difference between a clustering-based approach (*FindAllMarkers*) and the clustering-independent *singleCellHaystack*, we point out 7 example genes. For gene *Ctla2a* (Fig. 5A,H) both methods are in agreement; *Ctla2a* has very high expression in a subset of cells and is not detected in most other subsets.

**Figure 5:**
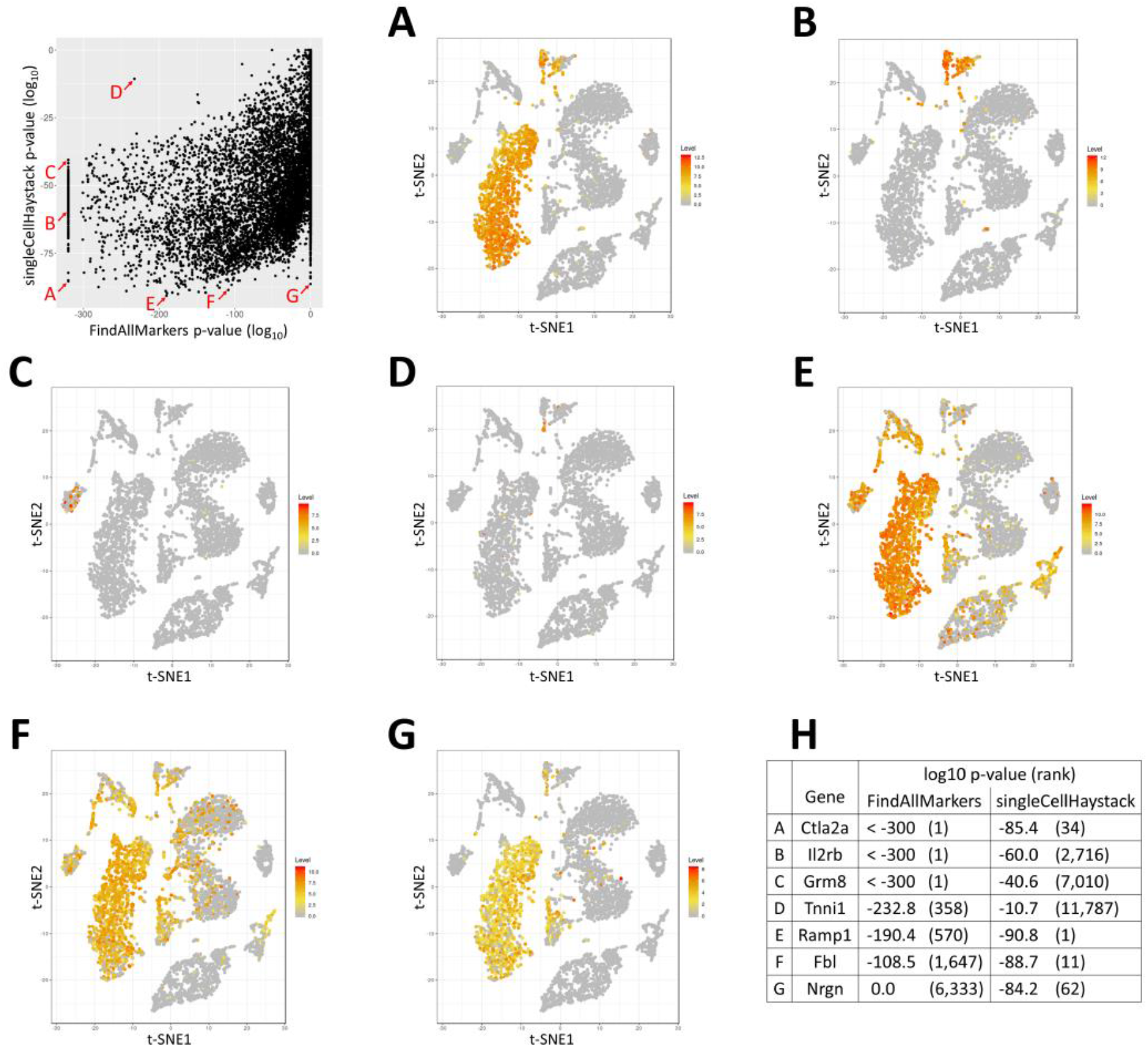
Comparison between *singleCellHaystack* and Seurat’s *FindAllMarkers* function on the Tabula Muris marrow tissue dataset. (top-left) Scatterplot of the p-values estimated by *FindAllMarkers* (X-axis) and *singleCellHaystack* (Y-axis) for all 13,756 genes in the dataset. 176 genes were given a p-value of 0 by *FindAllMarkers*, and are shown as p-value 1e-320. Expression patterns of indicated genes are shown in **(A-G)** and summarized in **(H)**.

Some genes are judged to have significant differential expression by *FindAllMarkers* but less so by *singleCellHaystack*. Gene *Il2rb* (Fig. 5B) is amongst the most significant genes reported by *FindAllMarkers* (p-values < 1e-300) but is not among the top ranked genes according to *singleCellHaystack*. The expression of this gene has a strong fit with the clusters underlying these results (Fig. S11). This trend continues with *Grm8* and *Tnni1*, which have high expression in a handful of cells within a single cluster (Fig. 5C,D and Fig. S11).

On the other hand, other genes are picked up by *singleCellHaystack*, but are not among the top-ranking genes according to *FindAllMarkers* (Fig. 5E,F,G). The most significant DEG according to *singleCellHaystack* is *Ramp1* (Fig. 5E, also shown in Fig. 2B). This gene is expressed across roughly half of the clusters as decided by *FindClusters* (Fig. S11), lowering its significance: *Ramp1* is ranked 570^th^ while *Tnni1* is ranked 358^th^ according to *FindAllMarkers*. A similar trend continues with *Fbl* and *Nrgn*, which have clear differential expression patterns but are not detected by *FindAllMarkers*.

These representative examples show that clustering-based approaches are likely to overestimate the significance of DEGs whose expression pattern fits very closely with a single cluster. These approaches are likely to miss DEGs whose expression is spread out over several clusters. On the other hand, *singleCellHaystack* can detect any pattern of differential expression, independently of clustering of cells. However, DEGs that are expressed in only low numbers of cells (ex: *Grm8*) might be missed.

## 4 Conclusions

*singleCellHaystack* is a generally applicable method for finding genes with biased expression patterns in multi-dimensional spaces. Although we have focused here on single-cell transcriptome data analysis, it is also applicable on large numbers of bulk assay samples, or on spatial transcriptome data. *singleCellHaystack* does not rely on clustering of cells, thus avoiding biases caused by the arbitrary clustering of cells. It can detect any non-random pattern of expression, and can be a useful tool for finding new marker genes. The *singleCellHaystack* R package includes additional functions for clustering and visualization of genes with interesting expression patterns.

As noted above, *singleCellHaystack* returns inflated p-values, because the input coordinates (PCs, t-SNE or UMAP coordinates) are dimensions which contain a large proportion of the variability in the original data. Clustering-based DEG prediction methods appear to suffer from this problem even more, because of their double use of gene expression data (for defining clusters and for DEG prediction) (13, 14). In future updates we hope to address this issue.

*singleCellHaystack* is implemented as an R package, available from https://github.com/alexisvdb/singleCellHaystack. The repository includes additional instructions for installation in R and example applications.

## Author contributions

A.V. conceived of the project and methodology and ran the analyses. A.V. and D.D. implemented the methods and wrote the manuscript.

## Acknowledgements

We thank the members of the Lab. of Systems Virology (Kyoto University), the Lab. of Functional Analysis *in silico* (Tokyo University), Dr. Yutaro Kumagai and Prof. Wataru Fujibuchi for helpful discussions and advice.

