## Supplementary Material for "*singleCellHaystack*: A clustering-independent method for finding differentially expressed genes in single-cell transcriptome data"

* To whom correspondence should be addressed.

**Availability and implementation:** https://github.com/alexisvdb/singleCellHaystack

**Contact:**

### Supplementary Material

### 1 *singleCellHaystack* methodology

The main function in *singleCellHaystack* is the `haystack` function. The input parameters to `haystack` are 1) the coordinates of cells in a multi-dimensional (≥2D) space (e.g. t-SNE, UMAP, or PC coordinates), and 2) data indicating for each gene in which cells it was detected and not detected. We refer to Supplementary Fig. S1 for an overview of the workflow.

#### Step 0: Optional normalization of input space

In this optional step, each dimension of the input coordinates is rescaled to mean 0 and standard deviation 1.

#### Step 1: Setting parameters

Steps 2-4 depend on using grid points for estimating the local density of cells using a Gaussian kernel, in function of their distance to the grid points. In the first step, `haystack_highD` decides the grid points and bandwidths for doing these calculations.

By default, 100 grid points are decided in a way that results in grid points being roughly uniformly spread over the subspace in which the cells are located. In other words, grid points should not be located close to each other, but should be proximal to the cells, and no cells should be distal from all grid points. In `haystack_highD`, the default way of deciding grid points is by running k-means clustering of the input cell coordinates, and to use the resulting 100 centroids as grid points. Note that the clustering of cells itself is not important. Another approach is to use seeding, as used in k-means++ clustering, in which initial centers are picked iteratively so that they are distal from other centers [1].

A bandwidth $h$ is decided as follows: for each cell, the distance to the closest grid point is calculated, and $h$ is defined as the median of those distances. Normalized distances between cells and grid points are subsequently defined as the Euclidean distances divided by the bandwidth $h$. The density contributions of each cell to each grid point along both axes is calculated as:

$$d_{cell,i} = e^{\left(-\frac{Dist_{cell,i}^{2}}{2}\right)}$$

where $Dist_{cell,i}$ is the normalized distance between $cell$ and grid point $i$.

##### Step 2: Estimating reference distribution $Q$

The density distribution of all cells in the multi-dimensional space is used as a reference distribution, $Q$. The density of cells at each grid point $i$ in the space is calculated as:

$$Q(i) = \sum_{cell} d_{cell,i}$$

After this, $Q$ is normalized to sum to unity.

##### Step 3: Estimating distributions $P(G = T)$) and $P(G = F)$)

The distribution of the cells in which gene $G$ is detected, $P(G = T)$, and those in which gene $G$ is not detected , $P(G = F)$, is estimated using the same bandwidth and grid points, as follows:

$$P(G = T, i) = \sum_{cell(G=T)} d_{cell,i}$$

and

$$P(G = F, i) = \sum_{cell(G=F)} d_{cell,i}$$

where $cell(G = T)$ and $cell(G = F)$ represent the subsets of cells in which $G$ is detected and not detected, respectively. Subsequently $P(G = T)$ and $P(G = F)$ are normalized to sum to unity.

##### Step 4: Estimating the Kullback-Leibler divergence of gene $G$, $D_{KL}(G)$

The divergence of the expression pattern of gene $G$, $D_{KL}(G)$, is calculated as follows:

$$D_{KL}(G) = \sum_{s\in\{T,F\}} \sum_{i} P(G = s, i) \log\left(\frac{P(G=s,i)}{Q(i)}\right)$$

If the cells in which $G$ is detected (and not detected) do not show a bias, and approximately follow the reference distribution $Q$, then $D_{KL}(G)$ is close to 0. As the discrepancy to the reference distribution $Q$ increases, the value of $D_{KL}(G)$ also increases.

##### Step 5: Estimating the significance of $D_{KL}(G)$

Finally, `haystack` evaluates the statistical significance of the $D_{KL}(G)$ values. We can not naively regard high $D_{KL}(G)$ values as significant, because there is a tendency for genes expressed in very few cells, and for genes expressed in almost all cells, to have a high $D_{KL}(G)$.

Instead, `haystack` evaluates the significance of observed $D_{KL}(G)$ values be comparing them to randomized data. First, let $c_G$ represent the number of cells in which gene $G$ is detected in the scRNA-seq data. Randomized genes are made that are expressed in $c$ cells. This is repeated many times, and the $D_{KL}(G)$ of these randomized genes are recorded. In practice, the values of $c$ are 200 values taken from the actual $c_G$ values of the data, evenly spread across the range of $c_G$ values, and 50 randomizations are done for each $c$ value. Since $c$ covers only 200 values (a subset of all unique $c_G$ values), this typically results in a much-reduced runtime. If there are less than 200 unique $c_G$ in the data, then all are used.

For each $c$ value, randomized $log_2(D_{KL}(G))$ follow an approximately normal distribution. We can therefore use the mean and standard deviation of randomized $log_2(D_{KL}(G))$ values to estimate a p-value of the actually observed $D_{KL}(G)$ values. The mean and standard deviation of randomized $log_2(D_{KL}(G))$ values are modeled as a function of $c$ using B-splines (using the `splines` R package). By default, 10 degrees of freedom are used for the splines, although this is lowered if there are few $c$ values.

Using the B-splines, expected means and standard deviations are predicted for each gene, in function of its detection count. P-values are then estimates using the `pnorm` function in R.

#### The *singleCellHaystack* advanced mode

In practice, in some cells more genes are detected than in others (for examples, see Fig. 4, left). The default reference distribution $Q$ does not take this into account, which can lead to genes being judged to be highly biased, even if they are merely detected in cells that have many detected genes.

To address this, the function `haystack` can take this into account by weighting the density contributions of cells by the number of genes detected in each cell:

$$d_{cell,i}^{advanced} = e^{\left(-\frac{Dist_{cell,i}{}^2}{2}\right)} \times g_{cell}$$

where $g_{cell}$ is the number of genes detected in $cell$. This assigns a larger influence to cells with many detected genes in the calculation of $Q$. Estimation of $P(G = T)$ and $P(G = F)$ is done as in the default mode.

A second difference is in the construction of the randomized "genes" (see Step 5 above). In the default mode randomized genes are simulated by randomly picking $c$ cells from all cells in the input data according to a uniform probabilities (i.e. each cell has the same chance of being selected). In the advanced mode, the probabilities reflect the number of genes detected in each cell (i.e. cells in which many genes are detected have a higher chance of being selected). As a result, randomized genes tend to follow the detection levels of the actual cells more closely.

The advanced mode can be activated by giving the vector of $g_{cell}$ values as input to the `use.advanced.sampling` parameter of the `haystack` function.

#### The `haystack_2D` function for 2-dimensional input

In addition to `haystack_highD`, which is aimed at >2D input, *singleCellHaystack* includes the function `haystack_2D`, which was specifically designed for 2D input coordinates. The general concept is the same. Below, we discuss differences in steps 1-4. Step 5 is the same as introduces above.

##### Step 1: Setting parameters

For `haystack_2D`, steps 2-4 depend on the mapping of 2D coordinates onto a grid, and estimating the local density of cells using a Gaussian kernel, in function of their distance to the grid points. In the first step, `haystack` decides the parameters for doing these calculations. Bandwidths $h_x$ and $h_y$ for the X and Y axes are decided using the rule-of-thumb of the `bandwidth.nrd` function (MASS package in R). The number of grid points is decided so that there are 25 grid points between the 10% and 90% percentile of coordinates along both axes. The grid is then further extended until it covers all points. The goal of this strategy is to reduce the influence of outliers on the definition of the grid.

The density contributions of each cell to each grid point along both axes is calculated as:

$d_{cell,i} = e^{\left(-\frac{\left(\frac{x_{cell}-x_i}{h_x}\right)^2}{2}\right)}$ for the X-axis grid points, and

$d_{cell,j} = e^{\left(-\frac{\left(\frac{y_{cell}-y_j}{h_y}\right)^2}{2}\right)}$ for the Y-axis grid points, where $x_{cell}$ and $y_{cell}$ represent the coordinates of each cell $cell$, $x_i$ and $y_j$ the coordinates of grid points of the X-axis and Y-axis, respectively.

##### Step 2: Estimating reference distribution $Q$

The density distribution of all cells in the 2D plot is used as a reference distribution, $Q$. The density at each grid point in the 2D plot is calculated as:

$$Q(i,j) = \sum_{cell} \sum_{i,j} d_{cell,i} \times d_{cell,j}$$

To each $Q(i,j)$ value, a small pseudo count is added, defined as the 1% percentile value of non-zero $Q(i,j)$ values. After this, $Q$ is normalized to sum to unity.

##### Step 3: Estimating distributions $P(G = T)$ and $P(G = F)$

The distribution of the cells in which gene $G$ is detected, $P(G = T)$, and those in which gene $G$ is not detected , $P(G = F)$, is estimated using the same bandwidths and grid points, as follows:

$$P(G = T, i, j) = \sum_{cell(G=T)} \sum_{i,j} d_{cell,i} \times d_{cell,j}$$

and

$$P(G = F, i, j) = \sum_{cell(G=F)} \sum_{i,j} d_{cell,i} \times d_{cell,j}$$

where $cell(G = T)$ and $cell(G = F)$ represent the subsets of cells in which $G$ is detected and not detected, respectively. The same pseudo count is added as for $Q$, and subsequently $P(G = T)$ and $P(G = F)$ are normalized to sum to unity.

##### Step 4: Estimating the Kullback-Leibler divergence of gene $G$, $D_{KL}(G)$

The divergence of the expression pattern of gene $G$, $D_{KL}(G)$, is calculated as follows:

$$D_{KL}(G) = \sum_{s \in \{T,F\}} \sum_{i,j} P(G = s, i, j) \log\left(\frac{P(G=s,i,j)}{Q(i,j)}\right)$$

##### Step 5: Estimating the significance of $D_{KL}(G)$

This step is the same as for `haystack_highD` (see above).

#### 2 scRNA-seq datasets and processing

Datasets were processed as described below. In many steps, we followed the recommendations described by Kobak and Berens [2].

##### Step 1: Data sources

- Tabula Muris data was download from https://tabula-muris.ds.czbiohub.org/ [3].
- Mouse Cell Atlas data file MCA_BatchRemove_dge.zip was downloaded from https://figshare.com/articles/MCA_DGE_Data/5435866. This data has been treated to reduce batch effects [4].
- The Nestorowa *et al*. dataset (processed read counts) was downloaded from GEO, accession number GSE81682 [5].

##### Step 2: Filtering of cells and genes

Library sizes of each cell were calculated by summing the number of reads or UMI (Universal Molecular Identifier) counts over all genes. Counts were then converted to counts per million counts (TPM). Genes were defined to be detected in a cell if their TPM was above a threshold TPM. The used thresholds were: 0 for the Tabula Muris microfluidic droplet data and the Mouse Cell Atlas data, and 32 for the Tabula Muris FACS-sorted data and the Nestorowa data.

In datasets with more than 20,000 cells, we randomly selected 20,000 cells. For the Tabula Muris microfluidic droplet data, we selected the 20,000 cells with the most detected genes.

We filtered out cells with fewer than 100 detected genes, and genes detected in fewer than 10 cells.

##### Step 3: Principal Component Analysis and t-SNE

We selected 1000 genes with large variance given their mean using dropout rates and mean TPM across non-zero counts, as described by Kobak and Berens [2]. The $\log_2$ TPM values of these 1000 genes (adding pseudocount of 1) were used as input for PCA, without scaling. Subsequently, t-SNE and UMAP was run on the first 5, 10, 15, 25, and 50 PCs. For t-SNE, we used the "Rtsne" package (version 0.15) [6]. t-SNE was run using perplexity 30, for a maximum of 500 iterations. For UMAP, we used the "umap" package (version 0.2.0.0) [7].

##### Step 4: *singleCellHaystack* analysis

*singleCellHaystack* was applied on the datasets using the `haystack` function, with as input 1) the coordinates of cells in a ≥2D space (2D t-SNE coordinates, 2D UMAP coordinates, or 5, 10, 15, 25, and 50 PCs) and 2) the detection data of each gene in each cell. This includes all genes that passed the filtering step (i.e. not only the 1000 genes used as input for PCA). `haystack` was run both using the default mode and the advanced mode which takes into general detection levels of genes (see main manuscript and explanation above). Run times were recorded for each run.

##### Step 5 Visualization of results

For the results returned by `haystack` by the default mode and the advanced mode, the following was done, separately:

- Genes with significantly biased expression patterns in the t-SNE plot were selected, using the function `show_result_haystack`. For this, the p-value threshold 1e-6 was used.
- Significantly biased genes were grouped into clusters by similarity of their expression pattern in the input space by hierarchical clustering. This was done using the function `hclust_haystack`. For the example application on the Mouse Cell Atlas Testis 1 dataset (Supplementary Fig. S8), the function `kmeans_haystack` for k-means clustering was used. In all cases the number of clusters was arbitrarily set to 5.
- The average distribution of the genes in each of the clusters was visualized using function `plot_gene_set_haystack` (see for example Supplementary Fig. 4).
- Within each cluster, the most significantly biased gene was plotted using `plot_gene_haystack` (see for example Fig. 1 and Supplementary Fig. S7 and S8).

#### Supplementary Figures

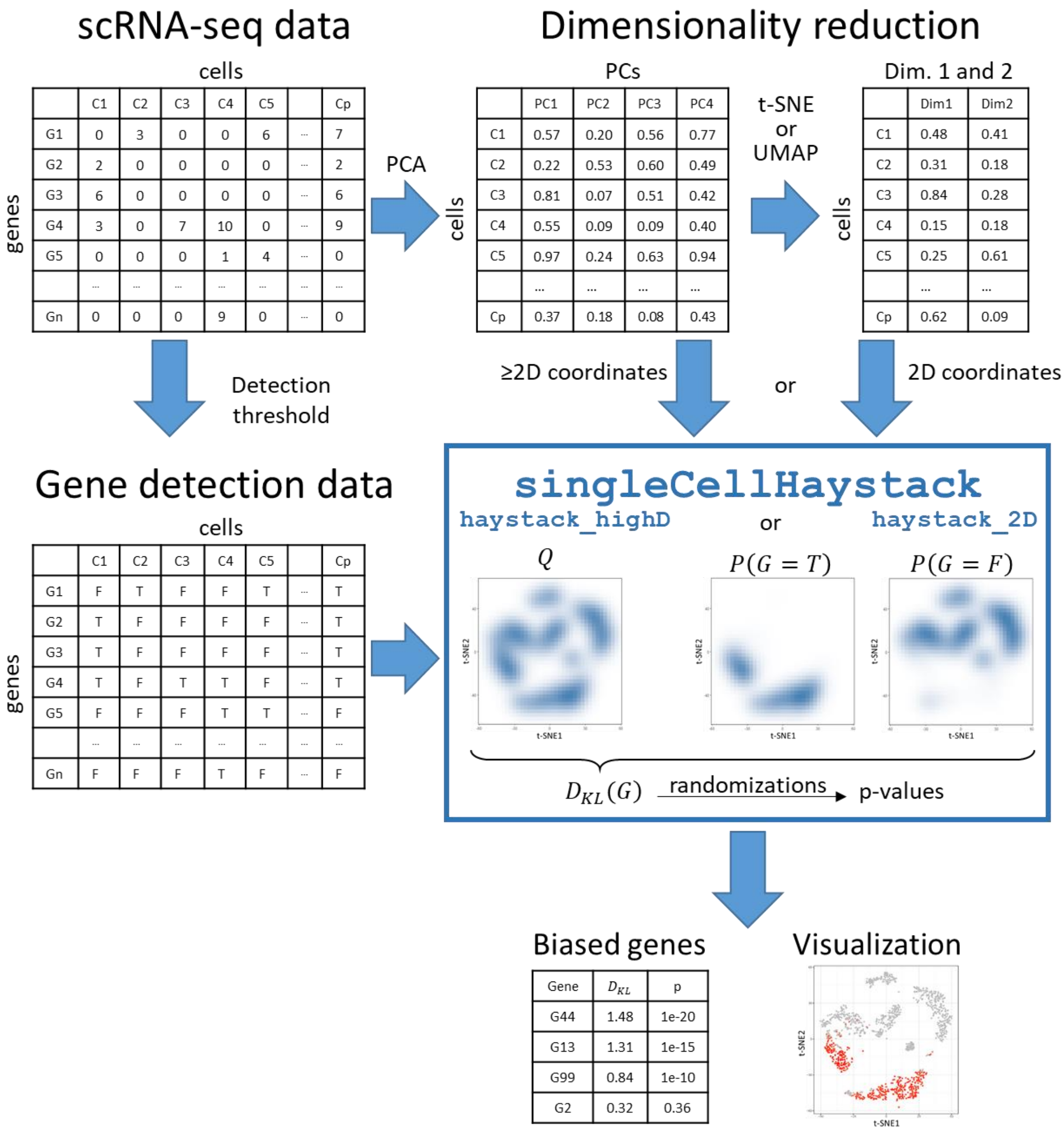

**Supplementary Figure S1: Overview of the *singleCellHaystack* workflow.** From the scRNA-seq data, cell coordinates in multi-dimensional space are obtained through PCA and (optionally) t-SNE or UMAP (or similar approaches). Read counts or UMI counts are converted to gene detection data (detected or not detected). The input to *singleCellHaystack* are the detection data, and multi-dimensional coordinates (`haystack_2D` for 2D coordinates and `haystack_highD` for ≥2D coordinates). The output is a list of all genes, their $D_{KL}$ and p-value. *singleCellHaystack* contains additional functions for visualization and clustering of genes according to their expression pattern in the multi-dimensional space.

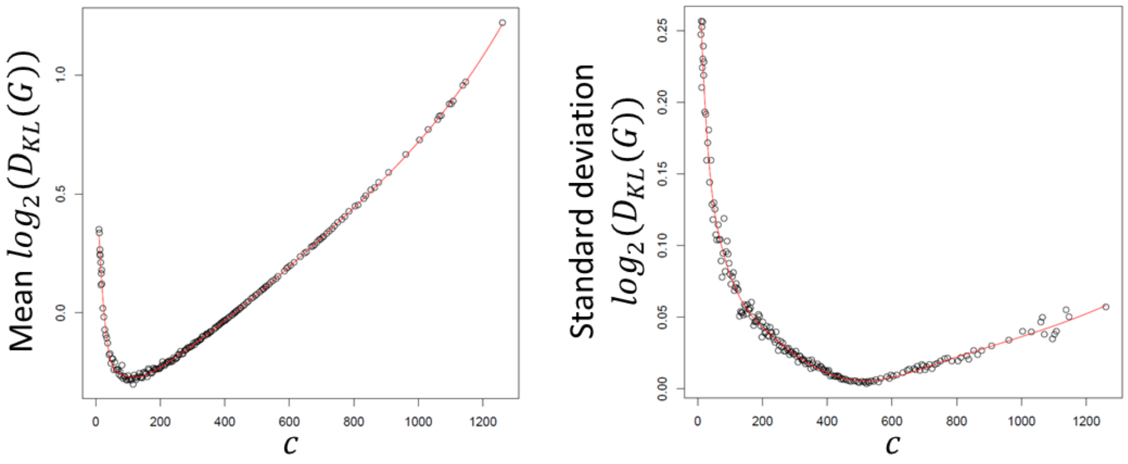

**Supplementary Figure S2: Example of trends of the mean and standard deviation of $D_{KL}(G)$.** The example shows the mean and standard deviation obtained from randomizations of the Mouse Cell Atlas muscle tissue dataset. Mean and standard deviations are shown for each value of $c$ (the number of cells in which a gene was detected) used in the randomizations. Fitted B-splines are shown in red.

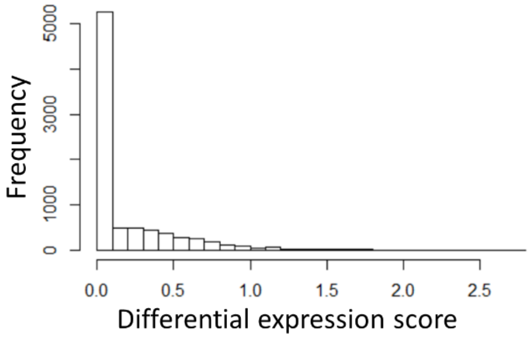

**Supplementary Figure S3: Histogram of the differential expression scores of genes in the artificial dataset generated using Splatter.** The 1,857 genes (out of 8,090) with a score > 0.3 were defined as DEGs.

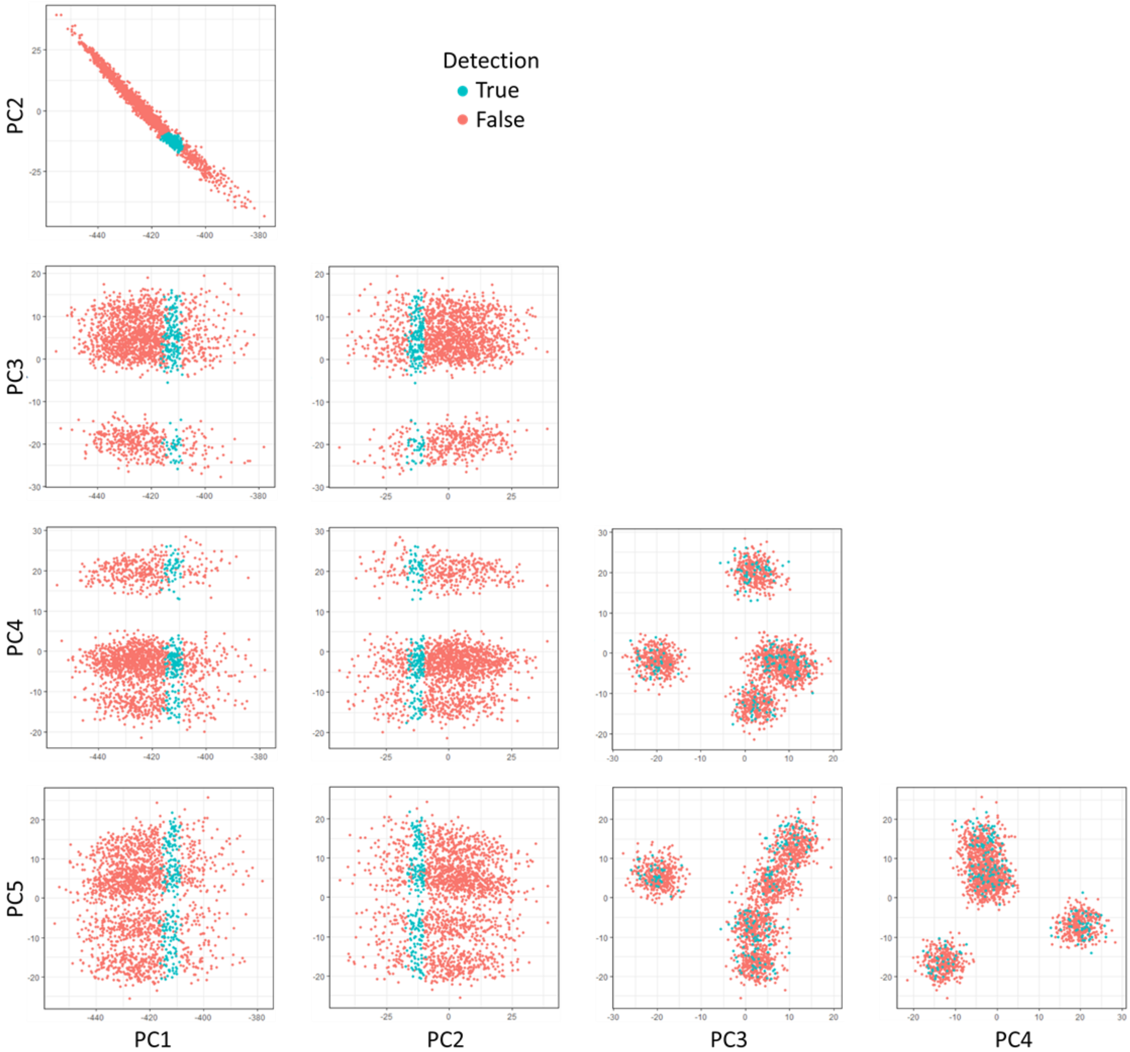

**Supplementary Figure S4: Expression data for the gene we manually added to the Splatter artificial dataset.** The scatterplots show pairwise PC coordinates of the cells colored by the detection (detected: cyan cells; not detected: orange cells) of the manually added gene. The gene is only detected in the 200 cells closest to cell No. 300 in the plane defined by the first and second PC. The corresponding t-SNE plot is shown in Fig. 1E.

**(next page) Supplementary Figure S5: Runtimes of *singleCellHaystack* and Seurat's *FindAllMarkers* function.** (A-B) Runtimes for `haystack_highD` on 50 PC input data, for the default mode (A) and the advanced mode (B). (C-D) Runtimes for `haystack_2D` on 2D t-SNE coordinates, for the default mode (C), and the advanced mode (D). (E-F) Runtimes of Seurat's *FindAllMarkers* function (the default Wilcoxon Rank Sum test), in function of number of cells in each dataset (E) and in function of the number of clusters of cells estimated in each dataset by the Seurat *FindClusters* function using 50 PC input data (F). Runtimes were measured on a Fujitsu Esprimo WD2/M (Intel® Core™ i7-4770 CPU, 3.40GHz).

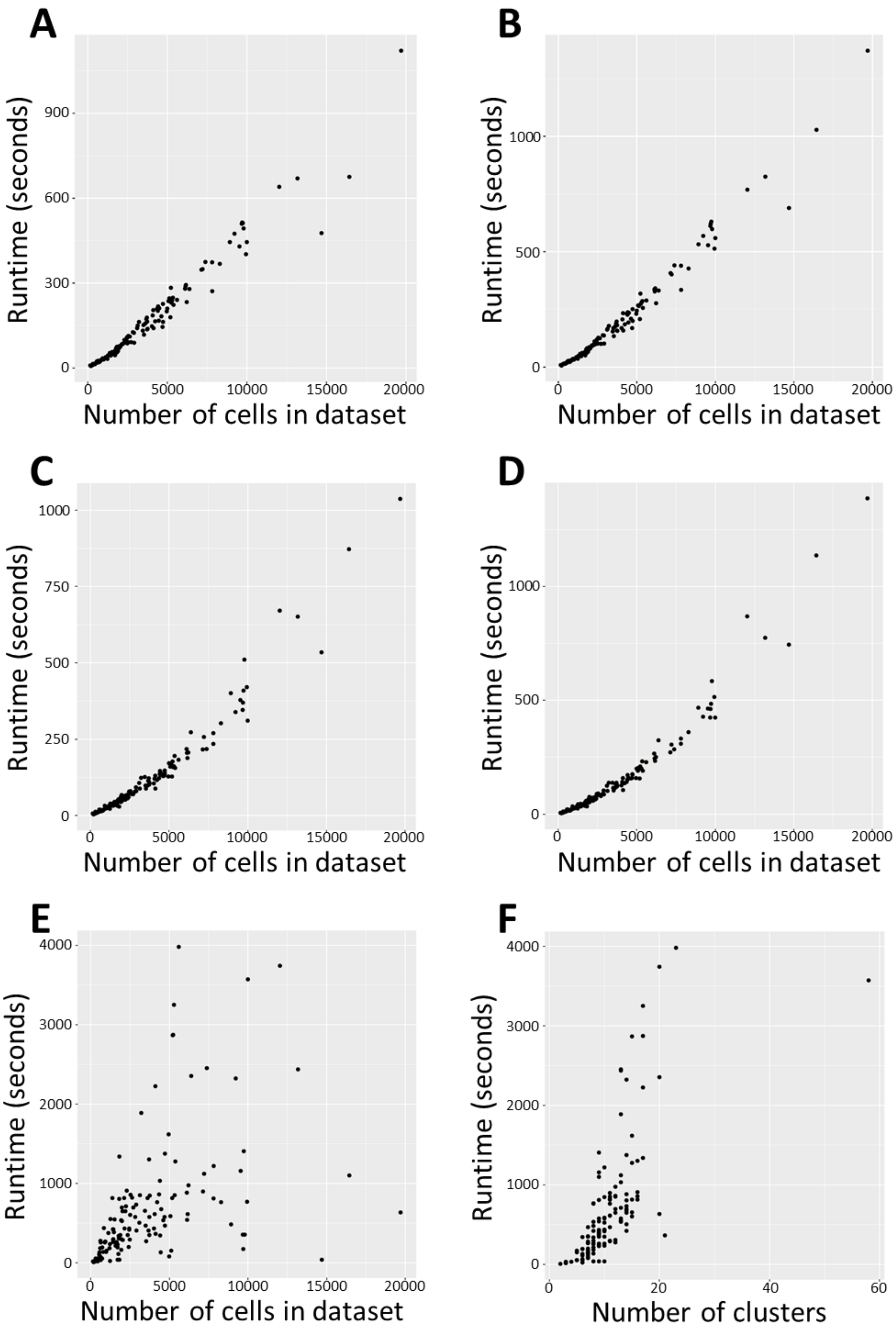
A
B
C
D
E
F
Runtime (seconds)
Number of cells in dataset
Number of clusters

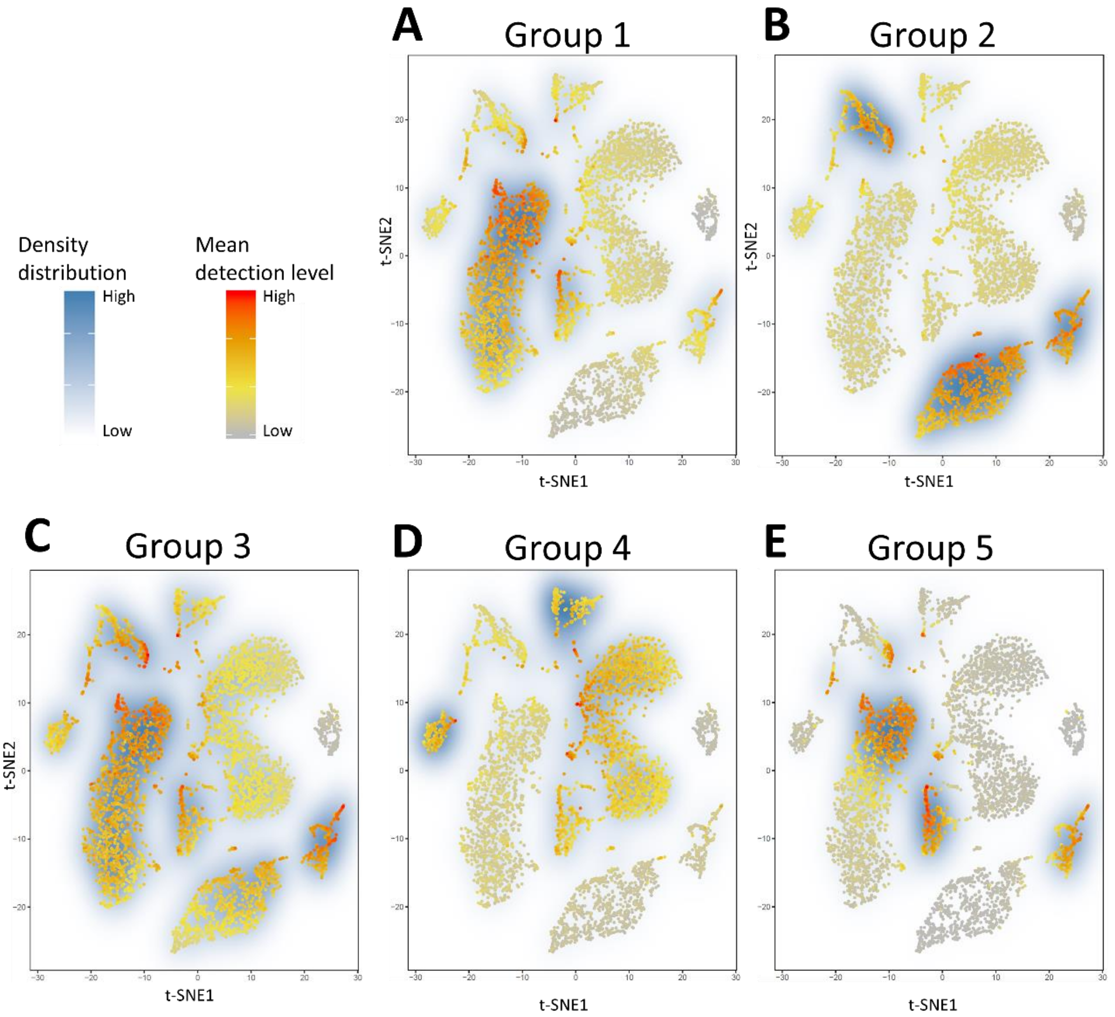

**Supplementary Figure S6: Clustering of biased genes in the Tabula Muris marrow tissue dataset.** Biased genes were grouped into 5 clusters using the `hclust_haystack` function, according to their density distribution in the input space (50 first PCs). (A-E) For each of the 5 resulting clusters, the mean detection level (the fraction of genes in the cluster detected in each cell) is shown (color scale from grey to red), as well as the averaged estimated density distribution of the genes in each cluster (color scale from white to blue). Genes shown in Figure 2 are the most significantly biased genes of each cluster.

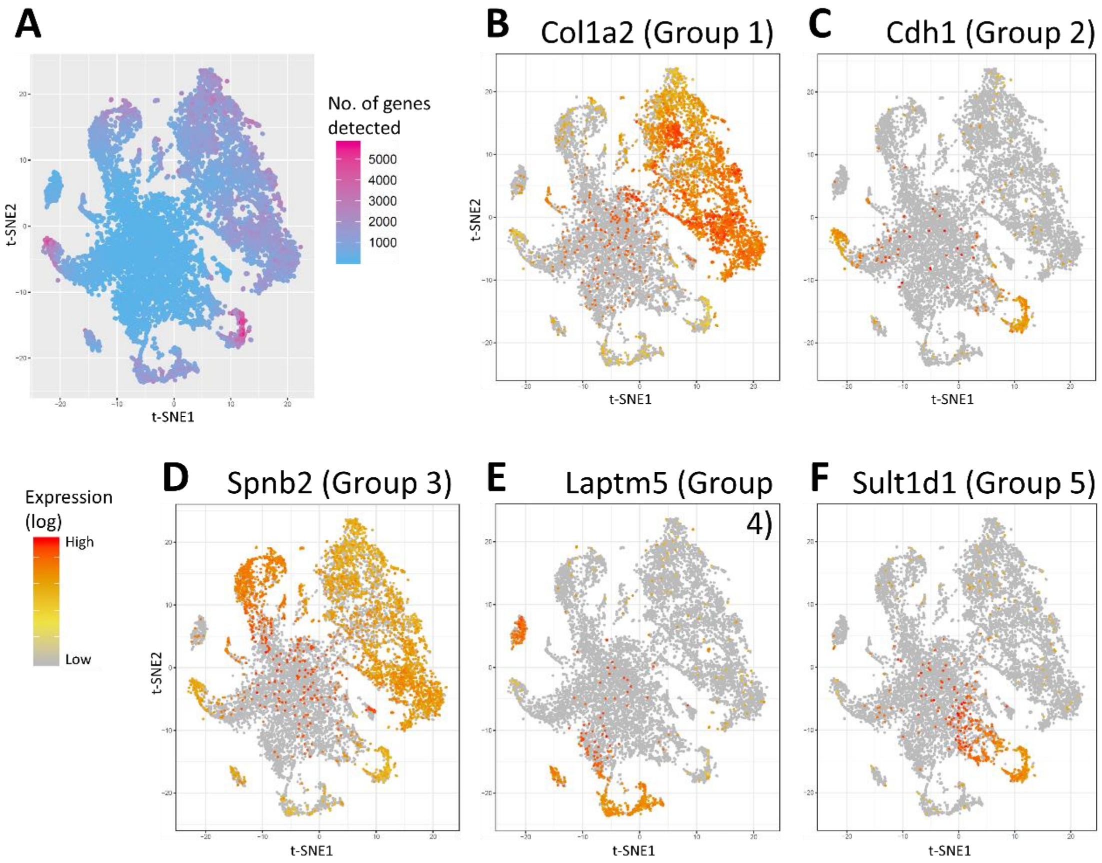

**Supplementary Figure S7: Application of *singleCellHaystack* on Tabula Muris (microfluidic droplet) Trachea (P8_14) tissue dataset. (A)** t-SNE plot of the 12,033 cells. The color scale shows the number of genes detected in each cell. **(B-F)** Expression pattern of five highly biased genes, representative of the five groups in which the genes were clustered.

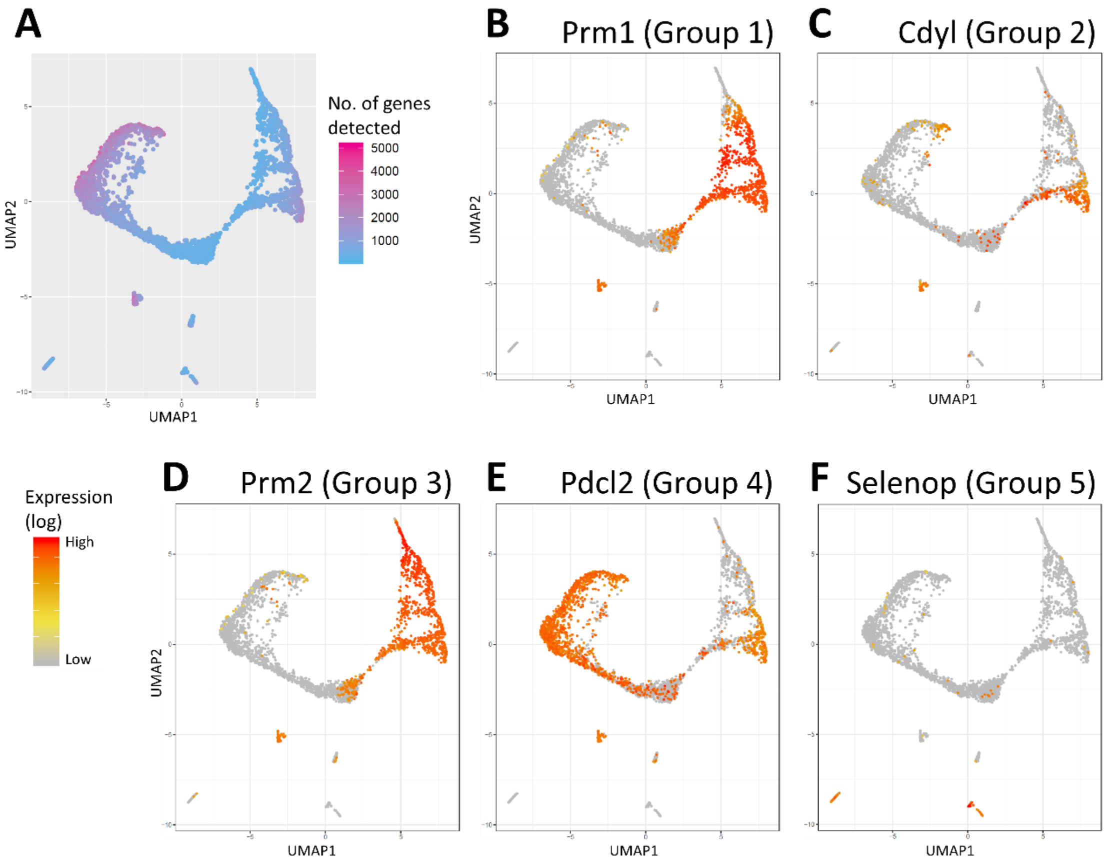

**Supplementary Figure S8: Application of *singleCellHaystack* on Mouse Cell Atlas testis 1 dataset. (A)** UMAP plot of the 3,217 cells. The color scale shows the number of genes detected in each cell. **(B-F)** Expression pattern of five highly biased genes, representative of the five groups in which the genes were clustered.

**(next page) Supplementary Figure S9: Consistency of *singleCellHaystack* results.** The distribution of ranks of genes in individual runs (Y-axis) is compared to the ranking using 5 PCs as input (based on mean rankings). **(A)** Rankings of individual runs on 5 PCs are in general consistent with the averaged ranking. **(B-D)** Consistency of results on 2D inputs: 2D t-SNE coordinates based on 50 PCs **(B)**, 2D UMAP coordinates based on 50 PCs **(C)**, and especially the first 2 PCs **(D)** result in higher variation of rankings. **(E-H)** Consistency of results on higher numbers of PCs as input: 10 PCs **(E)**, 15 PCs **(F)**, 25 PCs **(G)**, and 50 PCs **(H)** results in incrementally lower consistency (larger variation in the ranking of genes) with the averaged results based on 5 PCs.

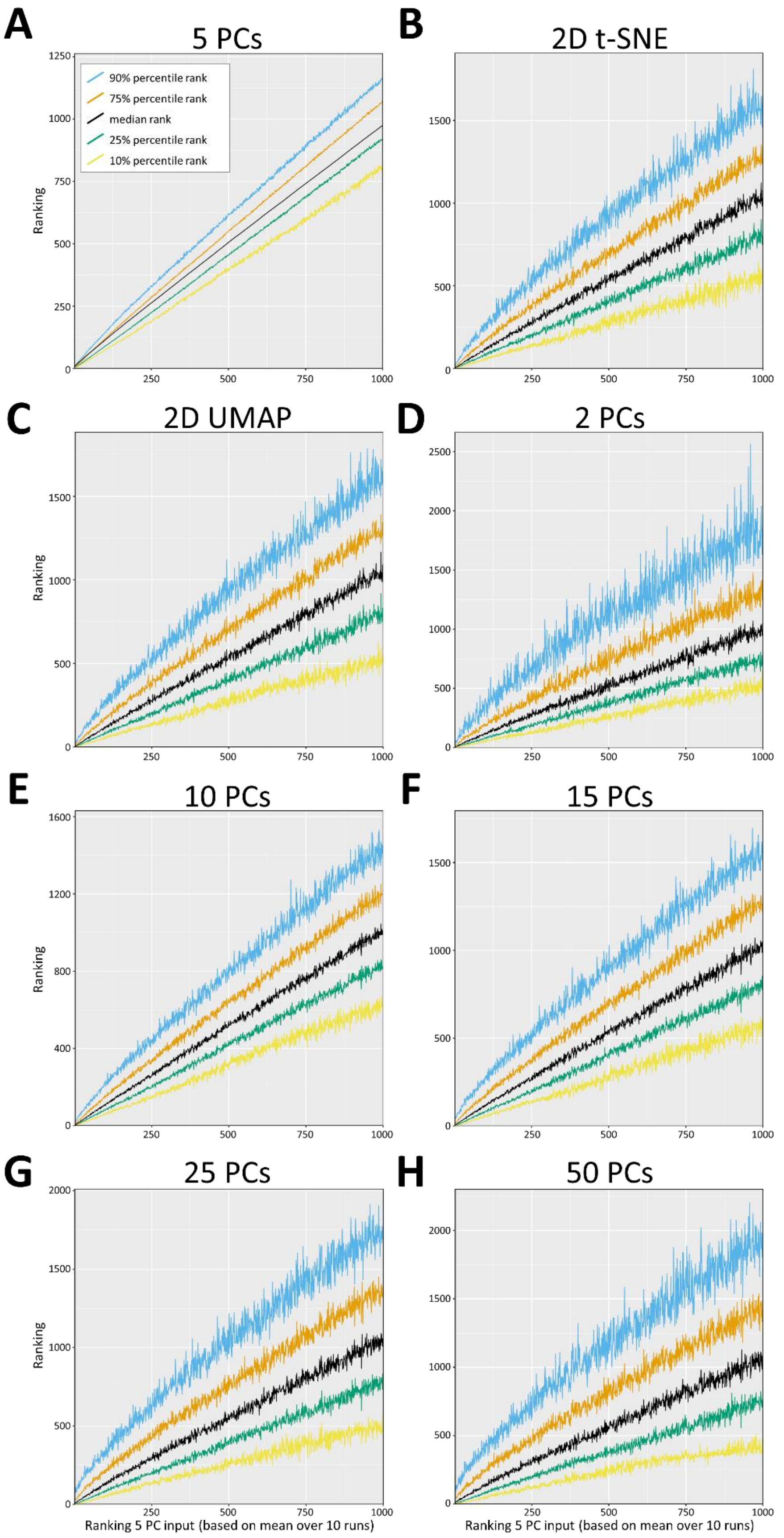
A
5 PCs
90% percentile rank
75% percentile rank
median rank
25% percentile rank
10% percentile rank
Ranking
B
2D t-SNE
C
2D UMAP
Ranking
D
2 PCs
E
10 PCs
Ranking
F
15 PCs
G
25 PCs
Ranking
H
50 PCs
Ranking 5 PC input (based on mean over 10 runs)
Ranking 5 PC input (based on mean over 10 runs)

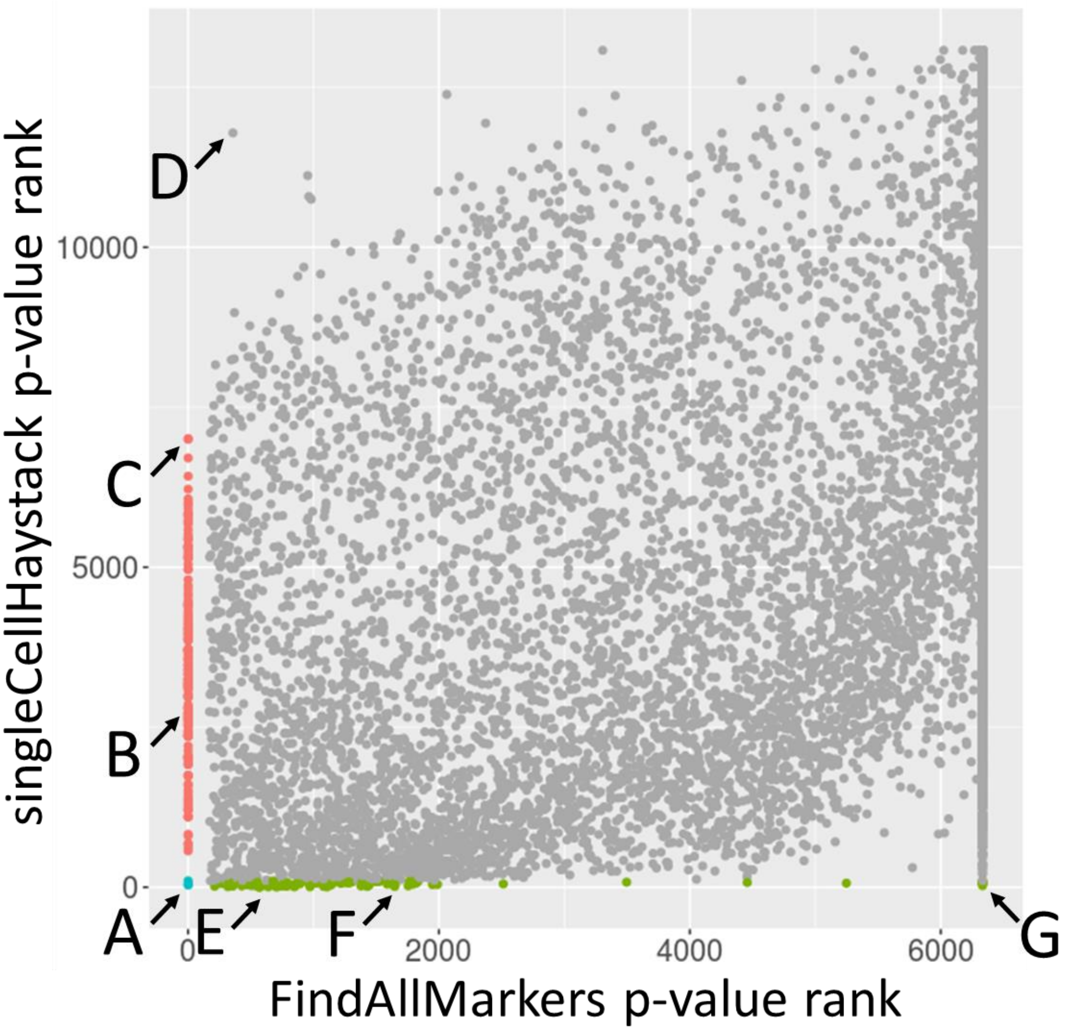

**Supplementary Figure S10: Comparison of ranking of genes by *singleCellHaystack* and Seurat's *FindAllMarkers* function on the Tabula Muris marrow tissue dataset.** Scatterplot of the ranks of p-values estimated by *FindAllMarkers* (X-axis) and *singleCellHaystack* (Y-axis) for all 13,756 genes in the dataset. Fig. 5 (top-left panel) in the main manuscript shows the scatterplot of p-values on which these rankings are based. Red: 176 genes with p-value of 0 by *FindAllMarkers*; Green: top 100 genes with highest significance according to *singleCellHaystack*. Cyan: three genes in the intersect of the above two sets of genes.

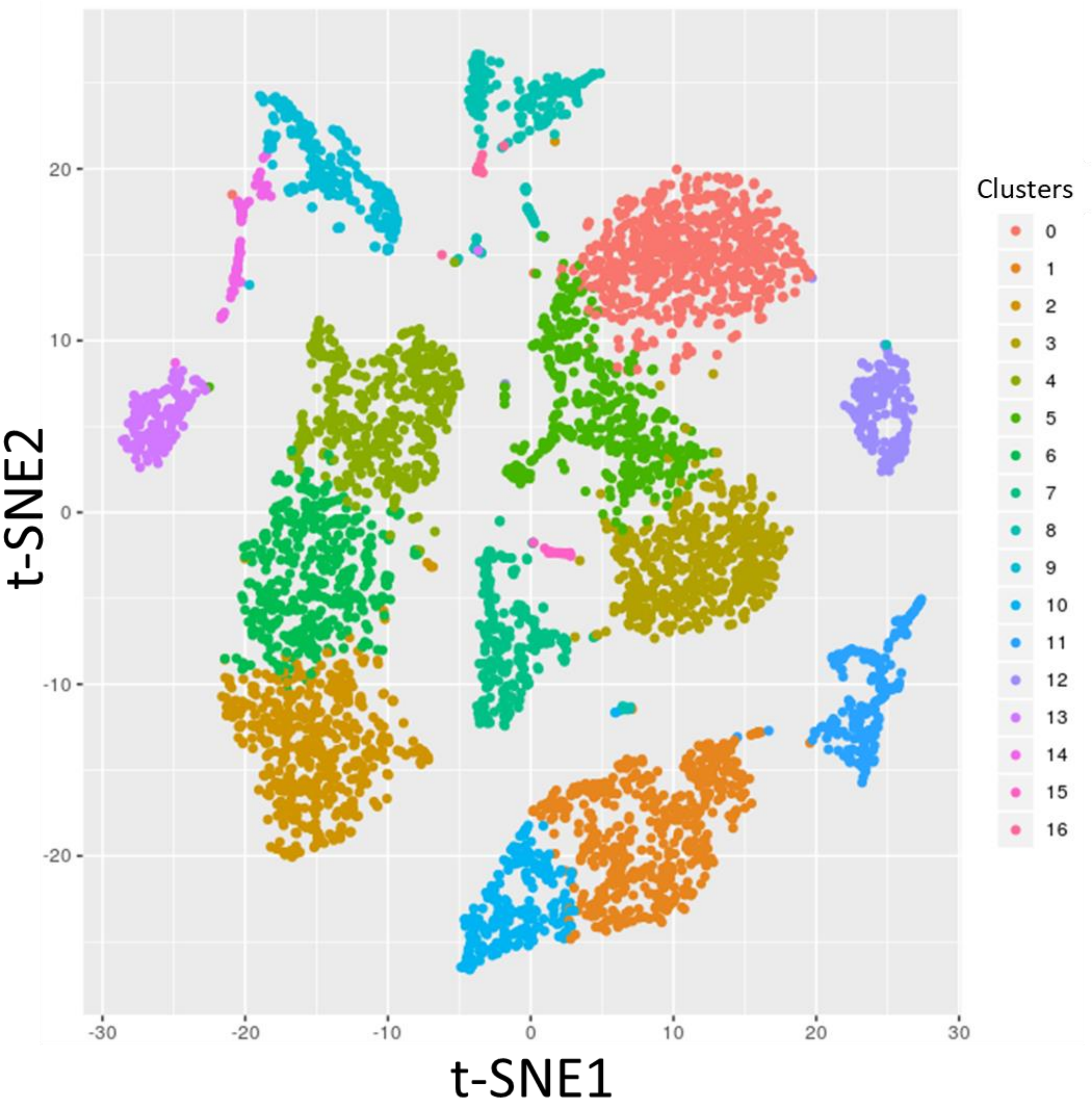

**Supplementary Figure S11: Clusters in the Tabula Muris marrow dataset decided by Seurat's FindClusters function.** Different colors represent different clusters of cells. There are 17 different clusters in total.
